# Computational simulation of ecological drift for generating functional minimal microbiomes identifies key experimental and biotic factors influencing success

**DOI:** 10.1101/2025.07.10.664178

**Authors:** Silvia Talavera-Marcos, Daniel Aguirre de Cárcer

## Abstract

We describe a top-down engineering approach that leverages ecological drift to generate *Minimal Microbiomes*; microbial consortia that are relatively simple, cohesive, and functionally complete. This process can be applied to any microbial ecosystem, provided that the target microbiome can be experimentally mimicked. Empirical support for this approach has emerged from multiple independent studies. Here, we use simulations across diverse scenarios, significantly varying niche structures and biotic interactions, to explore the experimental conditions and source microbiome characteristics that favor successful outcomes. Our results indicate that the effectiveness of this approach is constrained by several factors, and that perfect outcomes should not be routinely expected. Nevertheless, despite its drawbacks, this strategy remains a powerful tool for simplifying microbiomes and isolating key co-adapted populations, enabling the construction of low-diversity consortia that retain community function and present ecological cohesion.

## INTRODUCTION

In the natural environment, microorganisms are most frequently found as part of microbial communities; collections of potentially interacting populations that live together in the same space and time (1). These highly diverse communities play a crucial role in global biogeochemical cycles and form close associations with most multicellular organisms, significantly influencing their fitness.

The domestication of microbiomes is therefore a major goal in modern biology (2), with significant implications for agriculture, medicine, and biotechnology (3). Small, engineered synthetic consortia have been able to carry out functions such as bioremediation (4), manipulating plant phenotypes (5), or biofuel production (6), among others. Notwithstanding these stories of success, the bottom-up design of functional microbiomes remains challenging. This approach, selecting the right microbial populations to build a functional synthetic community, often requires a deeper understanding of specific microbial ecosystems, as well as microbial ecology and function in general, than we currently possess (7). Recognizing this limitation, Chang et al. introduced the idea that, rather than fighting unavoidable biological forces, an alternative top-down engineering approach could harness eco-evolutionary forces to create effective microbial consortia (2).

Attempts to remediate dysbiotic states in host-associated microbiomes (studies thus far overwhelmingly circumscribed to mammals and plants) using single bacterial populations have not been too successful. The scientific consensus nowadays is that complex consortia should be used instead of single strains, so that they can comprehensively replace the resident dysbiotic microbiota. The same is true for plant inoculants; single populations can seldom withstand the encounter with the rich reservoir represented by the native soil microorganisms. The key question then relates to how these complex communities should be constructed.

Many approaches have relied on rather observational information, constructing synthetic communities on the grounds of the taxonomic composition of “healthy” microbiomes using a library of isolates (a bottom-up approach). However, these approaches have so far provided limited success. For instance, in their seminal work, Ridaura et al (8) were able to modify the obese phenotype in mice using both co-housing with a lean mate and fecal transplantation experiments, but could not repeat the results using a specifically tailored consortium of 39 strains representative of the lean microbiota.

Microbial ecosystems most often consist of patches of strongly interacting dense microbial consortia (9), termed “local communities”. Within these microscale patches, the short distances between cells enable efficient diffusion-based exchange of metabolites, fostering strong biotic interactions that significantly shape the structure and dynamics of the community. Thus, populations that compete or antagonize one another may exclude each other within a single patch, but can still co-exist across the broader ecosystem when all patches (i.e., local communities) are considered collectively. Similarly, only co-adapted population pairs are able to inhabit the same patch, yet multiple such pairs can co-exist within a larger (macroscale) sample taken from the ecosystem.

Inevitably, the scale of most microbiological samples far exceeds that of the local community. This “Higher-scale sampling bias” (10), in which samples represent composites of multiple local communities, makes it extremely challenging to isolate co-adapted populations. In line with this rationale, the failure of Ridaura et al’s bottom-up approach may have resulted from a lack of co-adaptation among the selected strains. Therefore, how can we effectively bypass the Higher-scale sampling bias to isolate cohesive and functionally complete (i.e. local) microbial communities? An elegant yet untried solution is to actively manipulate ecological drift, defined as the stochastic changes in the relative abundance of populations in a community over time.

The cornerstone of the approach explored here, is a process (“the process” from now on) in which ecological drift is experimentally manipulated to access (i.e. isolate) local communities from complex microbial ecosystems. The underlying idea is that local communities represent functionally complete and cohesive microbial consortia. The overall idea of the process is relatively simple; microbial ecosystems consist of numerous microscale patches, within which individual local communities assemble. In a passage experiment, a sample (representing a macroscale portion of the ecosystem) is inoculated into a new, sterile environment. Bacteria are allowed to colonize this new environment, and the process is repeated multiple times. If the number of passages, colonization duration, and dilution rates between passages are properly calibrated, ecological drift should cause all local communities to eventually converge to a single community composition.

In most microbial ecosystems, the numerous microscale patches available to bacteria will fall into a limited number of categories, based on their abiotic conditions and(or) host-derived biotic environments in the case of host-associated microbiomes (e.g., normal epithelial patches vs. cecal crypts in the intestine). As a result, the process should yield homogeneous local community compositions for each type of patch. In this context, bacteria sampled from the resulting experimental ecosystem are likely to be co-adapted and capable of occupying all available niches within the ecosystem. Consequently, the resulting consortium could be considered functionally complete and cohesive. Naturally, the resulting microbiome will exhibit a significantly reduced richness (i.e., number of distinct microbial populations) compared to a typical sample from the same natural ecosystem, which would contain many different local community compositions across patch types. Therefore, generating a representative isolates library from the drift-simplified microbiome would entail substantially lower isolation costs compared to doing so from a natural, more complex sample.

Empirical support for the process has emerged in multiple independent studies. For instance, Goldford et al (11) investigated microbial community assembly using a single carbon and energy source. By cultivating complex microbial communities *ex situ*, derived from diverse natural environments, they observed that, for each compound, communities assembled into highly variable compositions at the finest phylogenetic resolution analyzed (i.e., exact sequence variants [ESVs] of the 16S rRNA gene). Despite this variability at the ESV level, communities consistently converged on the same family-level composition for each carbon source, regardless of their distinct environmental origins. In all passage experiments, the final communities were dominated by two populations, one from each of two bacterial families. However, different final communities harbored different inter-family pairs, indicating that all endpoint communities were composed of distinct homogenous local communities.

Second, Morella et al conducted a natural passage experiment on tomato leaf-associated microbiomes (12). As predicted by the drift-based process described above, they observed a general decline in community richness over successive passages. However, the authors attributed much of the lost diversity to the disappearance of transient or poorly adapted populations. This interpretation could have been influenced by their original experimental goal, which focused on assessing the relative contributions of environmental conditions and host genotype in shaping the resident microbiome. Most notably, the authors later performed a community coalescence experiment and found that the microbial communities from the final passage stages were resistant to invasion by the original bacterial communities. This outcome is consistent with the predictions of the process: homogenized local communities are functionally complete and cohesive, capable of occupying all ecological niches in the environment, and thus resistant to colonization by a diverse pool of incoming species.

The process, therefore, may serve as a valuable method for generating Minimal Microbiomes, microbial consortia that are relatively simple, cohesive, and functionally complete, from any microbial ecosystem, provided that the ecosystem can be experimentally mimicked and subjected to a passage experiment. The ability to obtain Minimal Microbiomes holds significant potential to drive the anticipated technological and economic advances in the field. These consortia could serve either as final products (with or without selection for specific community-level phenotypes) or as foundational materials for engineering genetically modified consortia, depending on the application area and, crucially, the intended end use.

A Minimal Microbiome may serve as the final product in applications that require only a stable, cohesive, and functionally complete microbial community; that is, a “healthy” microbiome. Such communities could be used to support host health in agricultural systems, aid in ecosystem restoration (e.g., following severe antibiotic treatment), or serve as standardized models for basic research.

Other anticipated applications will require the presence of specific community-level phenotypes in the original microbial sample. For example, disease-suppressive Minimal Microbiomes could be employed in agriculture, building on the well-established phenomenon whereby certain soils exhibit microbiota-driven resistance to specific plant pathogens (13). Similarly, microbiomes selected for host metabolic effects, such as weight-loss phenotypes to treat obesity or weight-gain phenotypes to address anorexia and support cancer patient recovery, could have therapeutic value in medicine. These types of phenotypes also hold relevance in livestock production, where antibiotics are still widely used in many countries to promote weight gain, despite their environmental impact. In the environmental and industrial sectors, Minimal Microbiomes with methanogenic capacity could be used to initiate anaerobic digesters for wastewater treatment, especially during start-up phases or after functional collapse, events that are not uncommon. Likewise, acetogenic Minimal Microbiomes could be developed to reduce methane emissions from rice paddies or ruminant livestock, both of which are major contributors to global greenhouse gas emissions. In each of these cases, the process could be used to generate Minimal Microbiomes starting from (meta)communities known to express the desired phenotype. These consortia would then be screened to identify those that retain the target functionality. The search for phenotype-associated consortia is often conducted using highly complex samples, making isolation and functional characterization difficult, or through combinatorial mixtures from isolate libraries, which often result in consortia with very few members, lacking both co-adaptation and functional completeness.

Finally, specific community-level phenotypes could also be achieved through the genetic modification of one or more individual populations within a Minimal Microbiome. For example, genetically modified bacteria (GMBs) have been proposed for use in the rhizosphere to release growth-promoting hormones or pathogen-inhibiting compounds. Similarly, GMBs are being developed for improved drug delivery in the human gut (14). These strategies could benefit substantially from being implemented within a Minimal Microbiome framework. Co-adapted microbial partners would help sustain the presence of the GMB in the environment, and the simplified structure of the community may facilitate the development of effective containment strategies. Moreover, this framework could enable the production of compounds that are difficult to synthesize within a single organism due to metabolic or genetic engineering constraints, by distributing the biosynthetic burden across the community. In agriculture, Minimal Microbiomes could also support the development of next-generation seed coatings (15) or other microbial formulations aimed at reducing agrochemical inputs while improving crop yield, nutritional quality, and resilience to biotic and abiotic stressors.

The overall goal of this study is to assess the feasibility of the proposed process for generating Minimal Microbiomes, and to explore under which experimental conditions and source microbiome characteristics might be successful. To this end, we developed a simulation framework that reproduces dilution–growth dynamics and predicts the final composition of microbial communities based on their initial characteristics and experimental parameters. We then used this toolkit to identify how those characteristics relate to the parameter settings that would be required to achieve fixation of non-redundant populations in a real-world passage experiment. To address this goal, we employed three simulation scenarios of increasing complexity, each progressively incorporating the following features: i) The presence of functionally redundant populations and stochastic growth. ii) The existence of functional groups with fixed relative abundances in the ecosystem and intra-group diversity. iii) The presence of interactions between populations belonging to different functional groups.

## METHODS

### Pipeline summary

The first step is to generate simulated initial communities with varying characteristics. The characteristics explored throughout this work include: the distribution of population abundances (uniform or log-normal), the number of distinct populations, the total community size (i.e., number of individuals), the number of functional groups and the assignment of populations to these groups, the relative abundance of each group (i.e., its niche size), and the presence or absence of biotic interactions.

The next step is to simulate the dilution–growth process with multiple replicates (i.e. trajectories) for each independent community. Several parameters must be specified, including the fixation threshold (i.e. the relative abundance above which a population is considered “fixed” within its functional group), the number of replicated trajectories, the number of dilution–growth cycles, and the dilution factor.

Each trajectory thus represents a single simulated community (i.e. the sum of all populations) undergoing dilution–growth cycles. During each cycle, the community is first diluted (i.e. subsampled) and then allowed to grow back to its original size. Growth is simulated through multiple iterations within each cycle. In each iteration, a fixed percentage of individuals is randomly selected to duplicate based on the relative abundance of each population. However, these probabilities can be modified by the presence of positive or negative interactions. In scenarios with defined functional groups (scenarios 2 and 3), random selection occurs within each functional group, and all groups grow until reaching their respective niche sizes. The final community composition at the end of each dilution–growth cycle is stored for downstream analysis.

The simulation results are analyzed to determine after how many dilution–growth cycles the fixation of a single population occurs within each functional group. This allows us to identify which dilution factor enables successful fixation within a defined number of cycles while maintaining the presence of all functional groups within the community (for scenario 2). To process and interpret the results, we define two key parameters: the success threshold and the fixation threshold. A community is considered successful when a “fixed” population appears in at least a given percentage of the simulated trajectories, where that percentage represents the success threshold. It is important to note that success does not require the same population to be fixed across all trajectories; rather, any population exceeding the fixation threshold within a sufficient proportion of trajectories qualifies the community as successful. On the other hand, the fixation threshold refers to the minimum relative abundance a population must reach in a given trajectory in order to be considered fixed. For example, let us consider a simulated dilution–growth experiment with a fixation threshold of 50% and a success threshold of 95%. For each community, 100 independent simulations (trajectories) are performed. In some trajectories, a single population may reach a relative abundance greater than 50% as early as cycle 20, while in others this may not occur until around cycle 25. However, if by cycle 30 95% of the trajectories show fixation of a single population (i.e. a relative abundance >50%), then we would consider that, under these fixation and success thresholds, the community reaches success at cycle 30.

The specific simulation scripts, along with the needed *dilgrowth* R package, are available in the GitHub repository github.com/silvtal/predicting_fixation, which also provides implementation details. The simulation workflow for each scenario is described in detail below.

### Scenario 1

For this opening exploration, we generated a set of initial communities by combining three parameters: richness (10, 100, or 1000 distinct populations), community size (10^4^ or 10^6^ individuals), and abundance distribution (uniform or log-normal). This resulted in 12 unique parameter combinations. For each unique combination, we constructed 30 replicate communities, yielding a total of 360 different initial communities. Each of the 360 initial communities was subjected to a series of simulated dilution-growth experiments, using one of 14 different dilution factors: 0.00025, 0.0004, 0.0005, 0.001, 0.0025, 0.004, 0.005, 0.008, 0.01, 0.025, 0.04, 0.05, 0.1, and 0.25. This resulted in a total of 5,040 distinct simulated experiments. Moreover, each experiment was independently replicated 100 times, yielding 504,000 individual simulation trajectories. Each trajectory consisted of 200 dilution-growth cycles, using a global growth rate of 1%. In the analysis of the results, a 95% success threshold was applied, along with two alternative fixation criteria set at 50% and 90%.

To quantify the effects observed during the graphical exploration of results, we modeled success by incorporating all relevant parameters using a random forest regressor implemented with the *party* package in R using 1000 trees. To this end, we applied a transformation to the target variable (success) to approximate a normal distribution using the Box-Cox transformation and thus help balance the data and ensure that the model performs with comparable accuracy across the full range of target values.

### Scenario 2

Here we established functional groups with a fixed relative abundance and stochastic growth of populations within each group. As in Scenario 1, we used 30 randomly generated communities for each community type. However, given the increased complexity of the simulations and based on the results from the previous scenario, we reduced the total number of parameter values explored. We therefore restricted our analysis to communities consisting of 10,000 individuals, with log-normal abundance distributions and initial richness levels of either 100 or 1,000. As a result, the number of distinct community types was reduced from twelve to two, defined solely by initial richness. Accordingly, the total number of initial communities decreased from 360 to 60. We analyzed nine different dilution factors instead of 14 (0.00025, 0.0005, 0.001, 0.0025, 0.005, 0.01, 0.025, 0.05, and 0.1) over 200 dilution-growth cycles. The success threshold remained at 95%, and the fixation threshold studied was 50%.

To incorporate the existence of functional groups, we used three or ten functional groups with either homogeneous or heterogeneous niche sizes (percentage of total community abundance assigned to each group), resulting in four different configurations (3 or 10 groups; homogeneous or heterogeneous niche sizes). The growth rate of each group was proportional to its niche size, with 1% growth per step distributed among the populations of each group according to their group’s niche size.

### Scenario 3

In this case, we explored the potential effect of inter-group interactions (i.e. between populations from different functional groups) on the overall success of the strategy, as well as on the fixation and extinction of functional groups. To do this, we reduced the number of communities studied to a single community from the previous scenario; one of the 30 communities with a log-normal abundance distribution and richness of 100. We analyzed the behavior of i) 3 groups with equal niche sizes and ii) 10 groups with heterogeneous niche sizes. For each community, 100 simulation runs were performed with 200 dilution-growth cycles at a single dilution factor: 0.1, the mildest dilution tested. This value was chosen because, in the previous scenario, success rates ranged from 0% to 95%, allowing for both potential increases and decreases in success rates, thereby revealing possible positive or negative effects of inter-group interactions.

The growth probability of each population was calculated as the sum of multiple components: its own relative abundance and the relative abundances of the populations it interacts with, weighted by the strength (positive or negative) of each interaction:

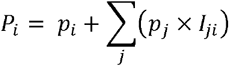

Where *p*□ is the relative abundance of population *i*, and *I*□□ represents the strength of the interaction from population *j* to population *i* (i.e., the corresponding value in the interaction matrix; see Supplementary Figure 1). We tested multiple types of interaction matrices, each defined by a unique combination of the following three parameters: i) sign (positive, negative, or both), ii) absolute value (1, 0.1, 0.01, 0.001, 0.0001), and iii) frequency (1%, 10%, 100%). The interaction frequency refers to the percentage of possible inter-group pairs, among all distinct pairs, that exhibit an interaction. All interactions were unidirectional. Each of the 100 simulations used a different randomly generated interaction matrix sharing one specific combination of these parameter values.

## RESULTS

Our initial simulation scenario was designed to explore the effect of the different experimental parameters (community richness, size and distribution, and dilution factor) on success, defined in this initial scenario as the fixation of a single population within the community (Figure 1, Supplementary Figure 2). First, we observed a strong effect of community size, with smaller communities reaching success more rapidly (Figure 1A-B). Second, there is a clear trend related to the dilution factor: within each community size, success is achieved earlier as dilution strength increases (Figure1C-D). In contrast, the abundance distribution appears to have little impact in most cases (Figure 1E-F) as did richness or diversity metrics (Supplementary Figure 2). We also noted that dilution factors above 0.1 or 0.25 (for 90% and 50% fixation thresholds, respectively) did not lead to successful outcomes, particularly for larger communities.

**Figure 1.**
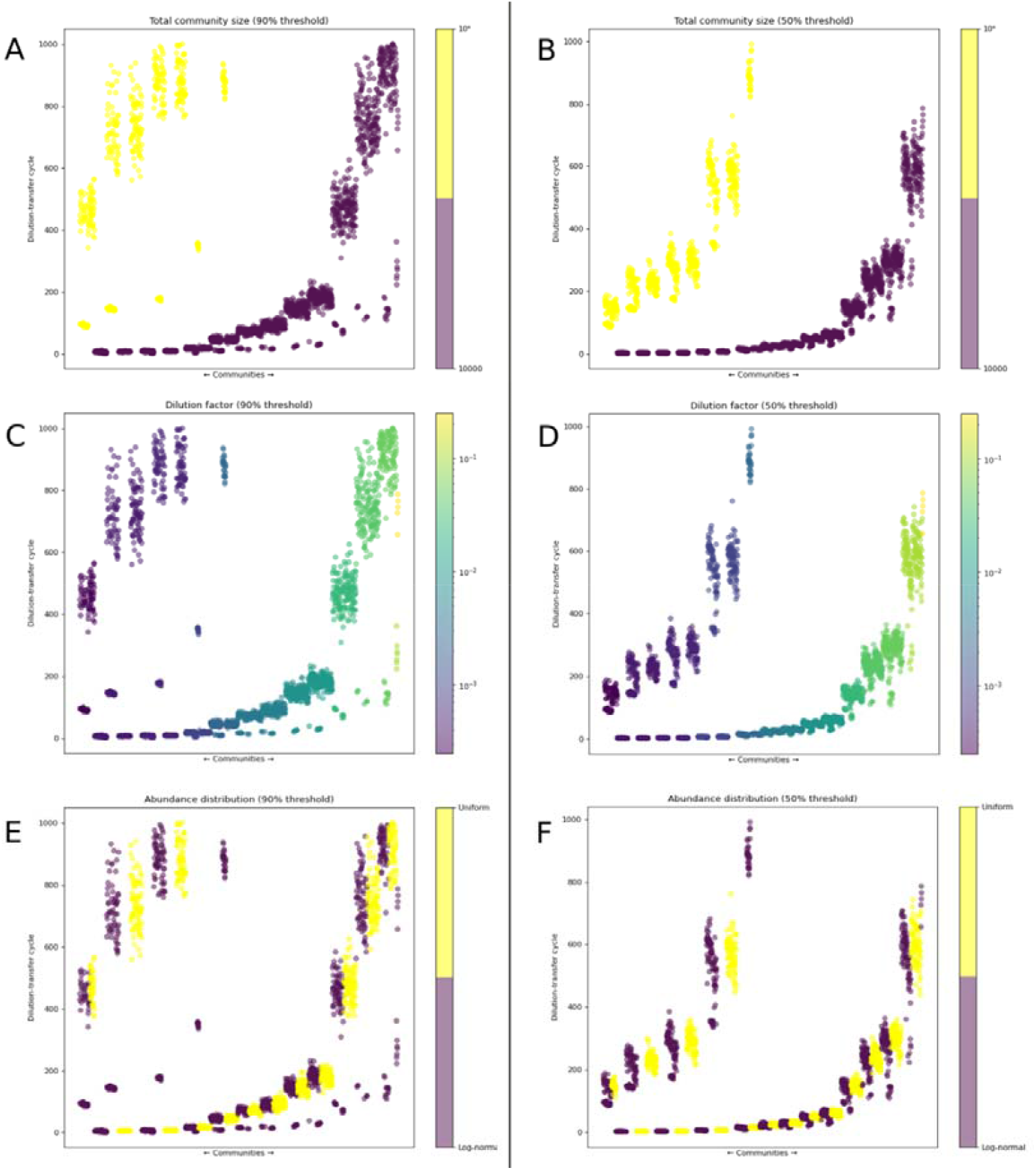
Success patterns of dilution-growth processes according to the simulation variables used. The Y axis indicates the dilution-growth cycle in which success occurred, while the X axis orders the different simulated communities by dilution factor, distribution, community size, and richness. In each plot, the simulations are colored according to one of the following variables, from top to bottom: community size (A, B), dilution factor (C, D), and population distribution (E, F). The plots show success according to two fixation thresholds: 90% in the first column and 50% in the second one. The success threshold is 95% in both cases.

Therefore, these dilution factors were excluded from subsequent analyses in this scenario, as well as from simulations in subsequent scenarios.

To quantify the effects observed during the graphical exploration, we constructed a random forest model. The model evaluates the importance of each variable by measuring the decrease in model accuracy when the values of that variable are randomly permuted. Variables whose permutation leads to a greater reduction in accuracy are considered more important. Consistent with the graphical observations, the model identified community size, dilution factor, and diversity metrics as the most influential variables, in that order (Table 1). However, recognizing that the inclusion of multiple diversity metrics introduces redundancy, we compared several new models, each including only one diversity measure, with or without richness as an additional variable. The results, based on the mean decrease in accuracy for each variable (Suppl. Tables 1 and 2) and the R^2^ values associated with each model (Suppl. Table 3), confirmed that the diversity metrics generally had a much weaker effect compared to community size and dilution factor. Among the diversity metrics, the Shannon diversity index exhibited marginally higher predictive power.

**Table 1.** Feature importance in the random forest model. Feature importance is calculated based on the mean decrease in accuracy. The values indicate how much the model’s accuracy drops when the variables are randomly permuted, for different fixation thresholds (50% and 90%).

| Fixation threshold | Community size | Dilution factor | Shannon diversity | Gini index | Pielou's evenness | Abundance distribution | Richness |
| --- | --- | --- | --- | --- | --- | --- | --- |
| 50% | 0.545 | 0.485 | 0.043 | 0.036 | 0.028 | 0.022 | 0.014 |
| 90% | 0.519 | 0.501 | 0.053 | 0.082 | 0.048 | 0.051 | 0.035 |

Next, we explored the success of the approach with simulated communities formed by 3 or 10 functional groups, each with its own fixed relative abundance and stochastic growth of intra-group populations. We observed that communities with 10 functional groups and heterogeneous relative abundances never achieved full success at any dilution factor, regardless of their richness (Supplementary Figures 3 and 4). This is expected since some groups have very low relative abundances (as low as 0.1%), making them highly vulnerable to extinction at each dilution step, even with milder dilution factors. In contrast, communities with 10 groups and homogeneous relative abundances showed fixation within a specific dilution range (0.005 to 0.1; Figures 2 and 3).

**Figure 2.**
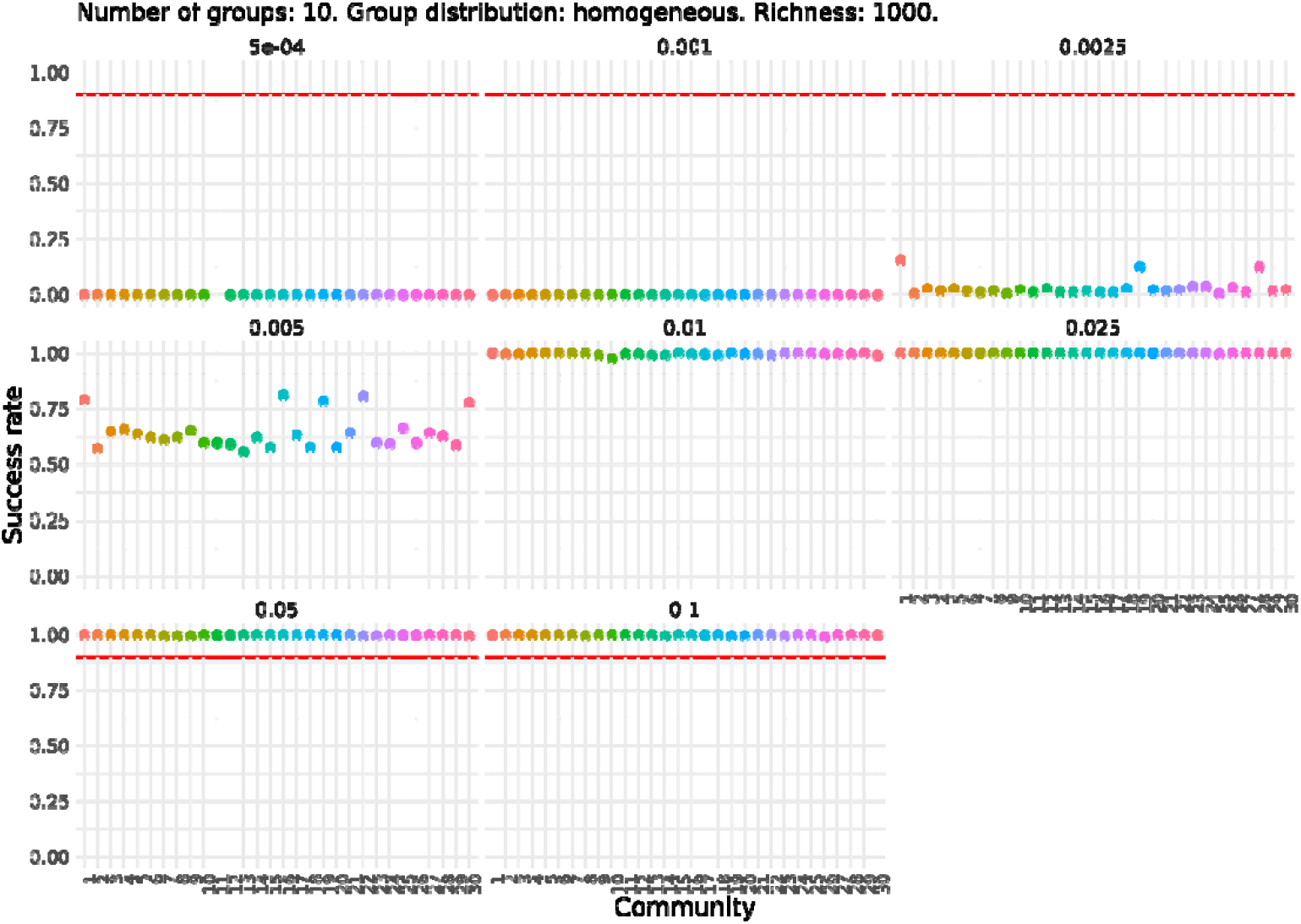
Success rate for each of the 30 simulated communities with 10 functional groups, homogeneous relative abundances, and a richness of 1000. Each plot corresponds to a different dilution factor. The Y axis indicates the proportion (from 0 to 1) of dilution-growth simulations in which total success occurs; that is, fixation in all functional groups. Each point represents one of the 30 communities. The missing point corresponds to a community that, after undergoing extinctions and a drop in total abundance, could no longer be sustained under the applied dilution factor.

**Figure 3.**
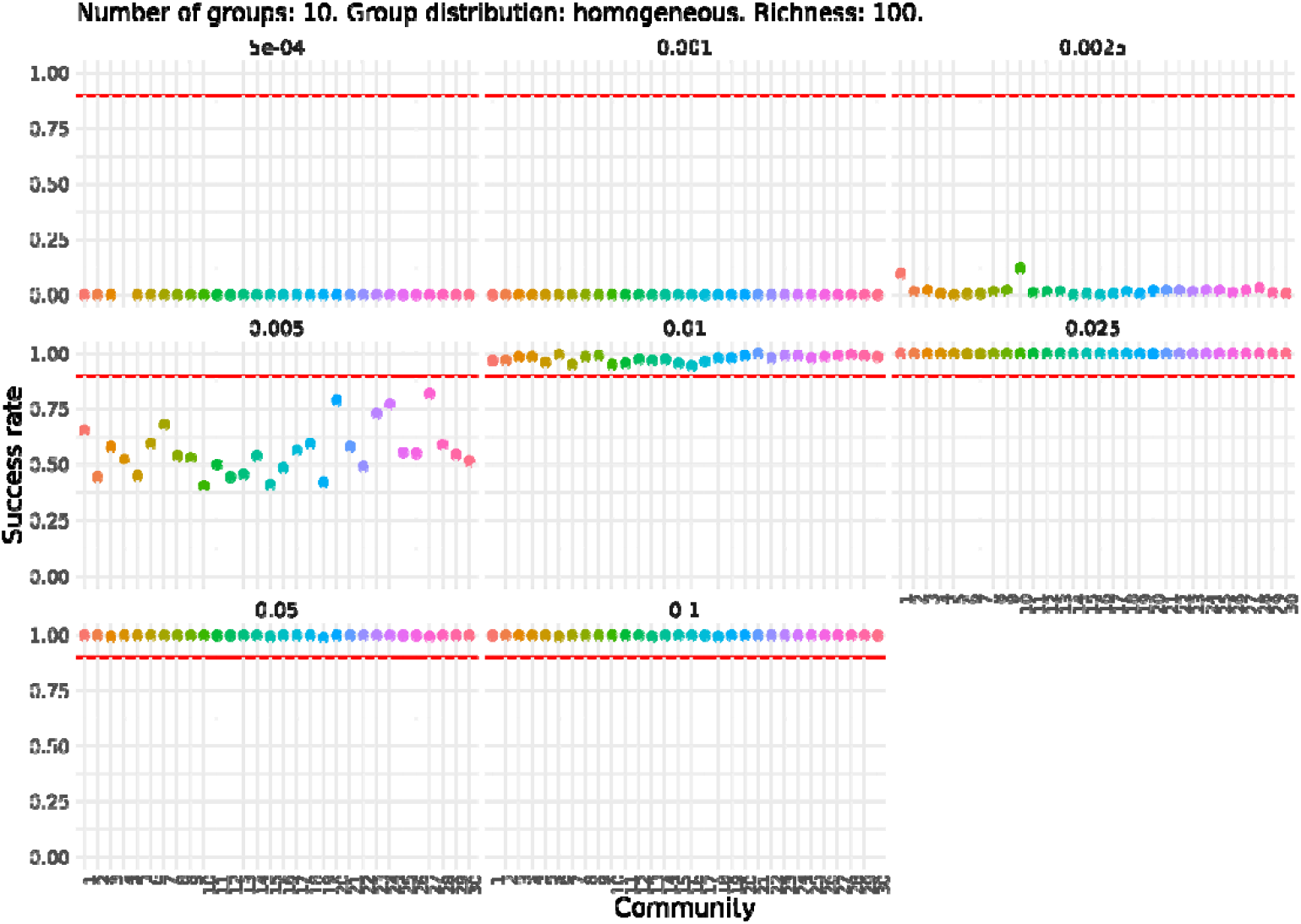
Success rate for each of the 30 simulated communities with 10 functional groups, homogeneous relative abundances, and a richness of 100. Each plot corresponds to a different dilution factor. The Y axis indicates the proportion (from 0 to 1) of dilution-growth simulations in which total success occurs; that is, fixation in all functional groups. Each point represents one of the 30 communities. The missing point corresponds to a community that, after experiencing extinctions and a decrease in total abundance, could no longer be sustained under the applied dilution factor.

For communities with 3 functional groups, regardless of niche size distribution or richness, we observed a dilution factor range where success increases with dilution intensity. In communities with equal niche sizes, results were similar for both richness values (Supplementary Figures 5 and 6), between dilution factors 0.0025 and 0.05 all communities reached fixation of the three groups. However, at higher dilution intensities (0.001 and 0.0005) and the lowest dilution intensity (0.1), fixation occurred but with success rates below the threshold. The rise and then fall in success rates as dilution decreases suggests that weak dilutions may prevent larger groups from achieving fixation. For communities with variable niche sizes (Figures 4 and 5), the dilution range with success rates above 95% was narrower (0.005–0.05), likely because less abundant groups were more prone to extinction.

**Figure 4.**
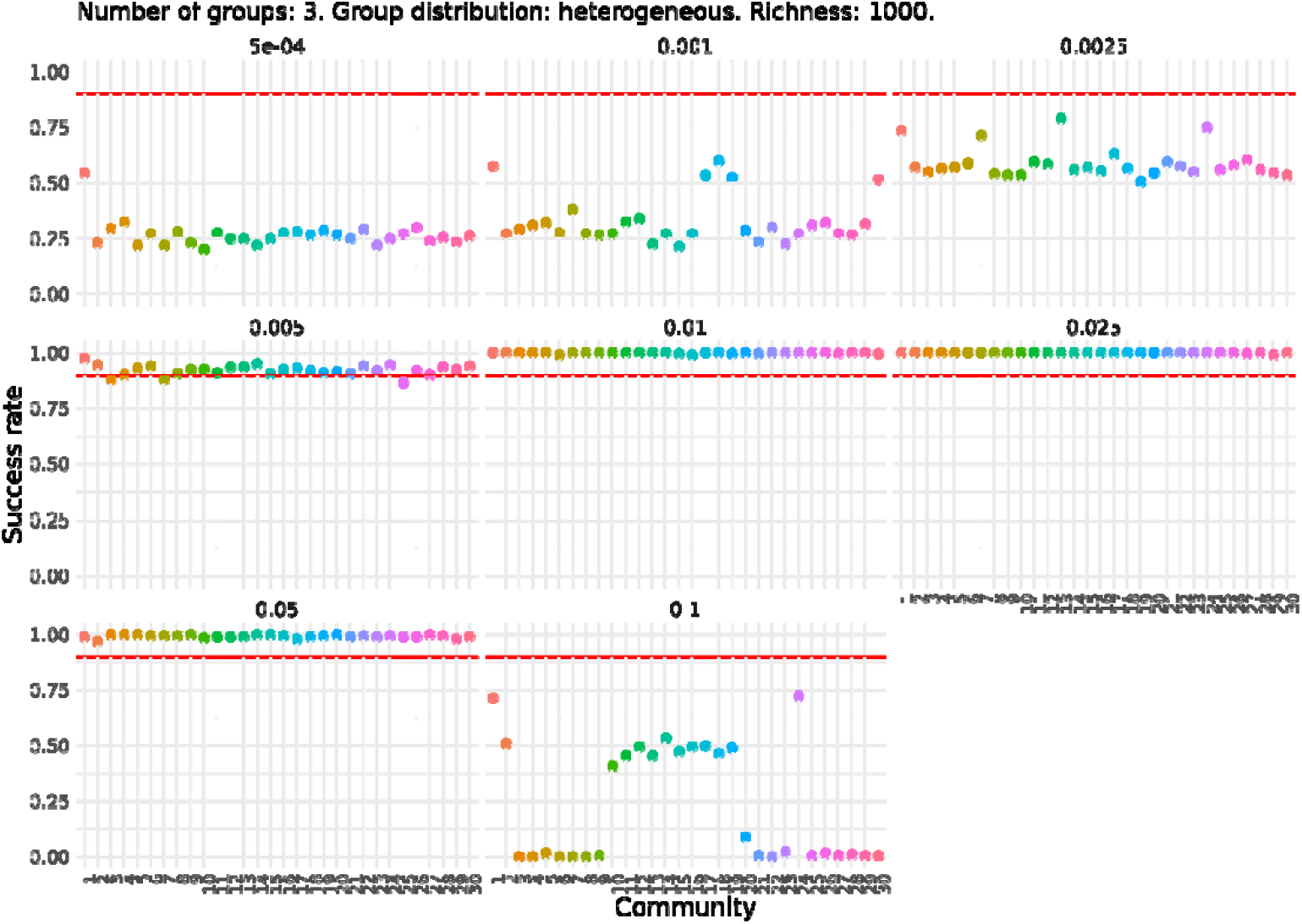
Success rate for each of the 30 simulated communities with 3 functional groups, heterogeneous relative abundances, and a richness of 1000. Each plot corresponds to a different dilution factor. The Y axis indicates the proportion (from 0 to 1) of dilution-growth simulations in which total success occurs; that is, fixation in all functional groups. Each point represents one of the 30 communities.

**Figure 5.**
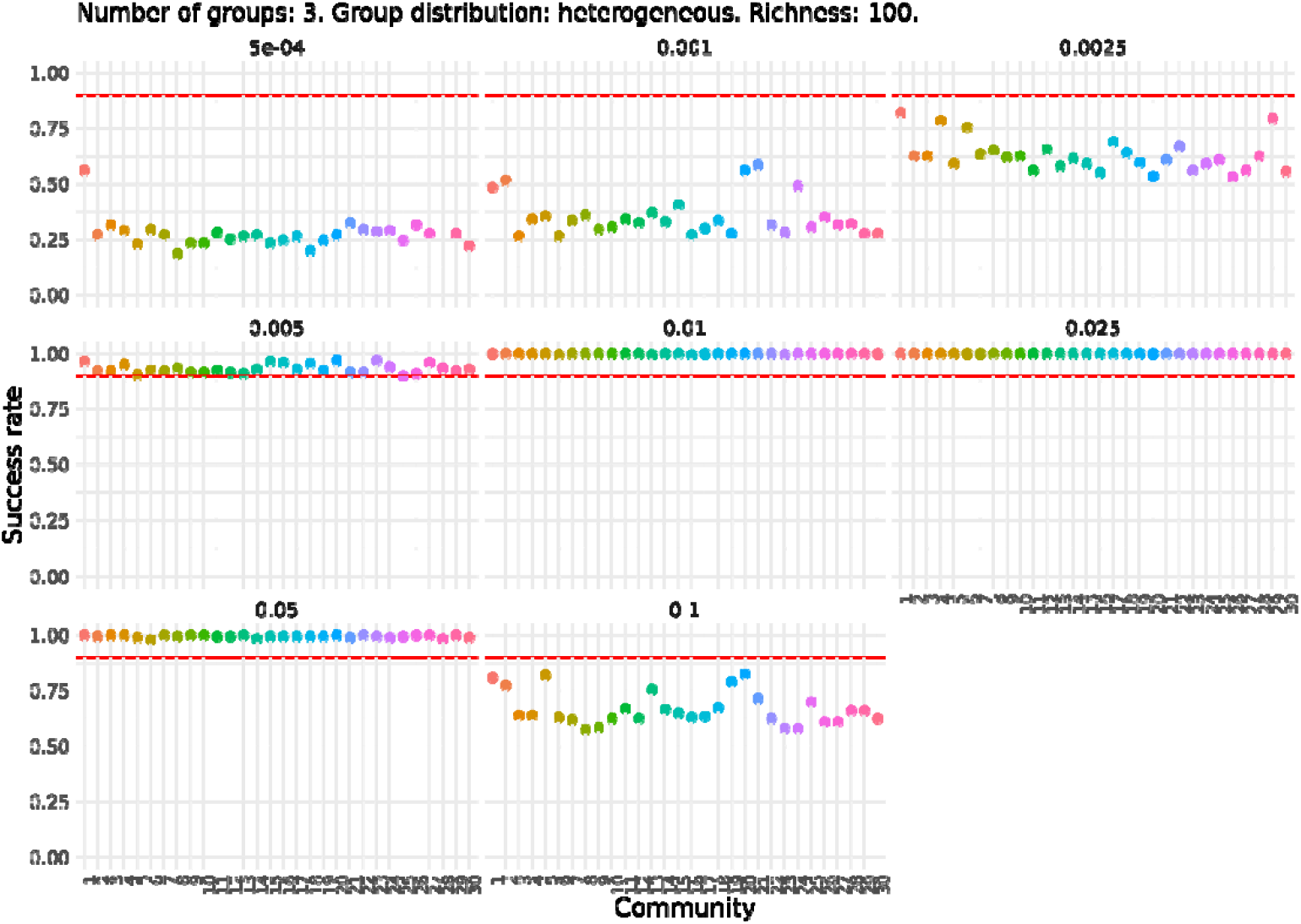
Success rate for each of the 30 simulated communities with 3 functional groups, heterogeneous relative abundances, and a richness of 100. Each plot corresponds to a different dilution factor. The Y axis indicates the proportion (from 0 to 1) of dilution-growth simulations in which total success occurs; that is, fixation in all functional groups. Each point represents one of the 30 communities. The missing point corresponds to a community that, after experiencing extinctions and a decrease in total abundance, could no longer be sustained under the applied dilution factor.

Next, we examined the fixation rates of each individual group (Figure 6, Figure 7, Supplementary Figures 7 and 8). As expected, in communities with homogeneous niche sizes, fixation of one group generally coincides with fixation of all groups. In communities with heterogeneous niche sizes, less intense dilution led to lower fixation rates in the larger groups compared to smaller ones. However, the two smallest groups were exceptions, showing near-zero fixation rates across all tested dilution factors, likely due to their higher extinction risk.

**Figure 6.**
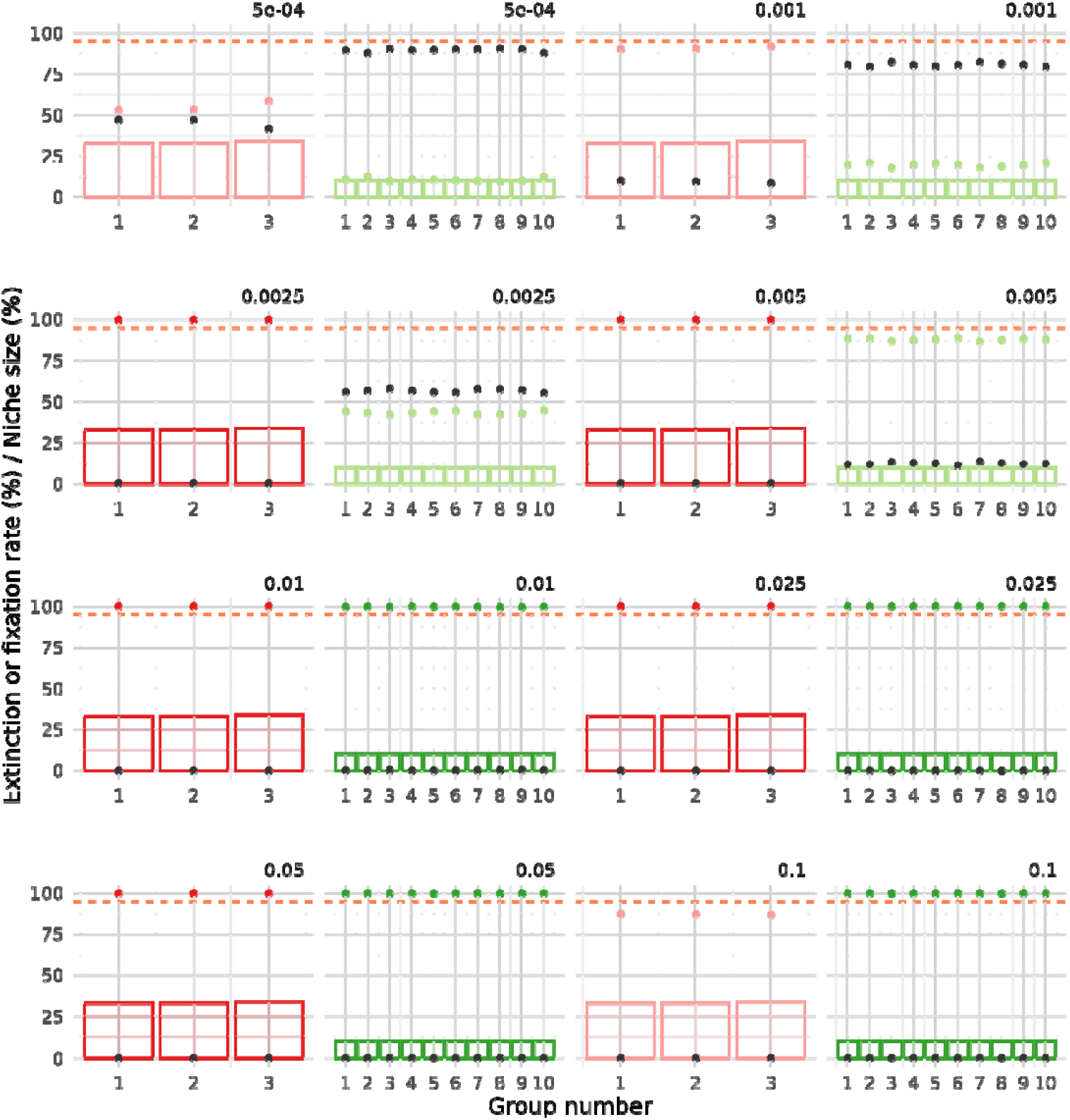
Fixation and extinction rate by functional group in communities with homogeneous niche abundances. Results are shown for communities with a richness of 100 and 3 or 10 groups (fixation in red and green, respectively, extinction in dark grey). Fixation for groups that exceed the 95% success threshold is highlighted in a darker shade. Bars represent the niche size associated with each functional group. Fixation is measured as the percentage of simulations in which at least one population from the group reaches fixation. Extinction is measured as the percentage of simulations in which no population from the group reaches fixation.

**Figure 7.**
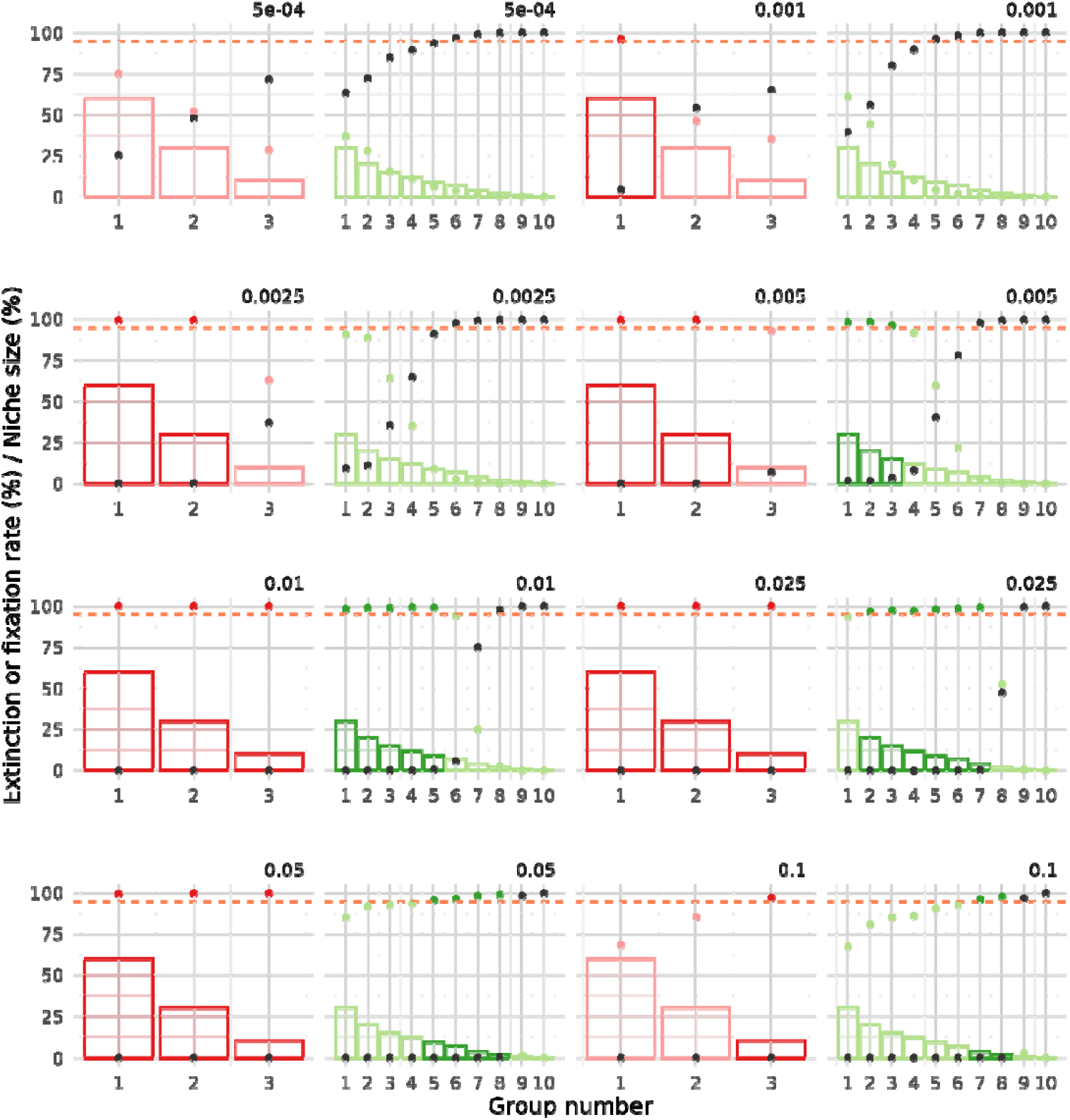
Fixation and extinction rate by functional group in communities with heterogeneous niche abundances. Results are shown for communities with a richness of 100 and 3 or 10 groups (fixation in red and green, respectively, extinction in dark grey). Fixation for groups that exceed the 95% success threshold is highlighted in a darker shade. Bars represent the niche size associated with each functional group. Fixation is measured as the percentage of simulations in which at least one population from the group reaches fixation. Extinction is measured as the percentage of simulations in which no population from the group reaches fixation.

For communities with 3 groups, success was achieved for all groups at dilution factors between 0.01 and 0.05, regardless of richness. In communities with 10 groups, partial success was observed: fixation occurred in one or more groups but not simultaneously in all. For both richness levels (100 and 1,000), the optimal dilution factors were 0.01 and 0.025, allowing up to 5 or 6 groups to exceed the 95% success threshold. To further investigate the observed effects, we examined the process from the perspective of group extinction (Figure 6, Figure 7, Supplementary Figures 7 and 8). We found that the failure to achieve fixation in certain groups was mainly due to their extinction caused by excessively strong dilution factors, rather than an insufficient number of cycles. In summary, the results highlight a delicate balance between extinction and fixation among groups with both larger and smaller niche sizes.

Finally, we explored the potential effect of inter-group interactions (i.e. between populations from different functional groups) on the overall success of the strategy, as well as on the fixation and extinction of functional groups. For the community with 3 groups of equal niche size, we observed that infrequent interactions led to an increase in fixation rates, regardless of interactions sign (Figure 8). The increase in fixation was more pronounced with positive interactions (either solely positive or mixed signs) at the two highest interaction strength values. This effect can be interpreted as an advantage for a limited number of populations over others. When interactions were exclusively negative, fixation rates improved as interaction frequency increased to 10% or 100%. One possible explanation is that these negative interactions promoted the extinction of some populations, thereby facilitating fixation of unaffected populations. However, this pattern did not hold when negative interactions had both an absolute strength of 1 and a 100% frequency; in this case, simulated growth was effectively suppressed. Finally, introducing strong and frequent positive interactions, whether alone or combined with negative ones, resulted in a sharp decline in fixation rates. This suggests a stabilization of a high number of intra-group populations, preventing fixation of any single population.

**Figure 8.**
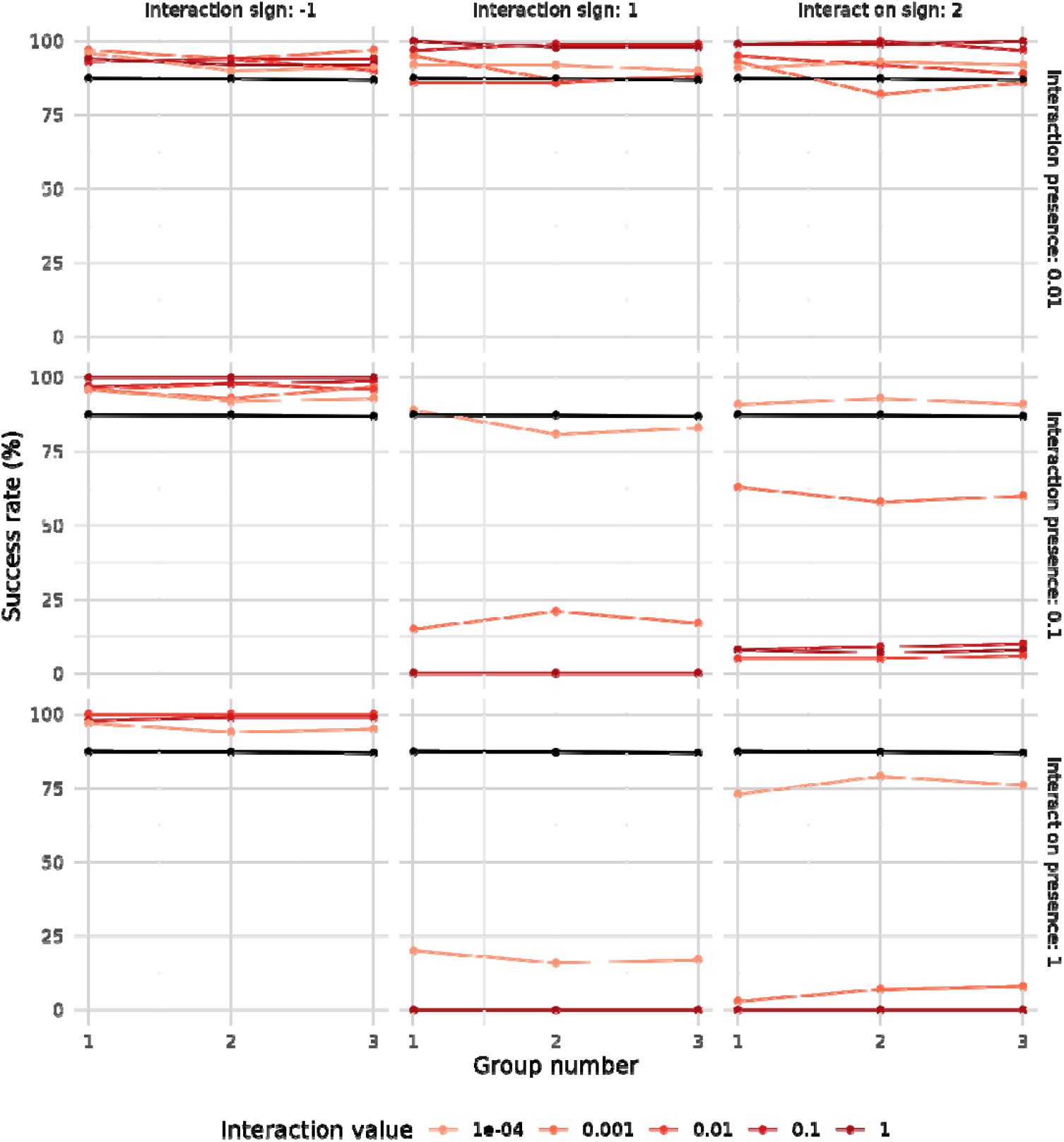
Success rate by functional group in communities with 3 groups (homogeneous group sizes, richness 100) following simulations under different interaction scenarios. Results without interactions are shown in black. Results for varying interaction strengths are displayed in different colors, indicating both the presence and the sign of the interaction. A dilution factor of 0.1 was used in all simulations.

For the community with 10 functional groups of heterogeneous niche sizes, we again observed that negative interactions generally promoted higher fixation rates across all groups (Figure 9). This effect was consistent across various combinations of interaction strength and frequency, though some exceptions were noted. In contrast, negative interactions with 100% frequency tended to suppress overall community growth. Once again, introducing positive interactions, either alone or in combination with negative ones, produced similar outcomes to infrequent negative interactions, as long as they remained rare. In other words, infrequent interactions consistently led to increased fixation rates, regardless of sign or absolute strength. On the other hand, simulations with positive interactions at medium to high frequencies generally resulted in a decrease in fixation rates, and this decrease was proportional to both interaction frequency and strength. The only exceptions occurred when the strength of positive interactions was low (0.0001) and either i) the interaction frequency was 10%, or ii) the frequency was 100% but included negative interactions as well. This suggests that positive interactions must remain limited for fixation rates to increase, mirroring the pattern observed in the 3-group communities.

**Figure 9.**
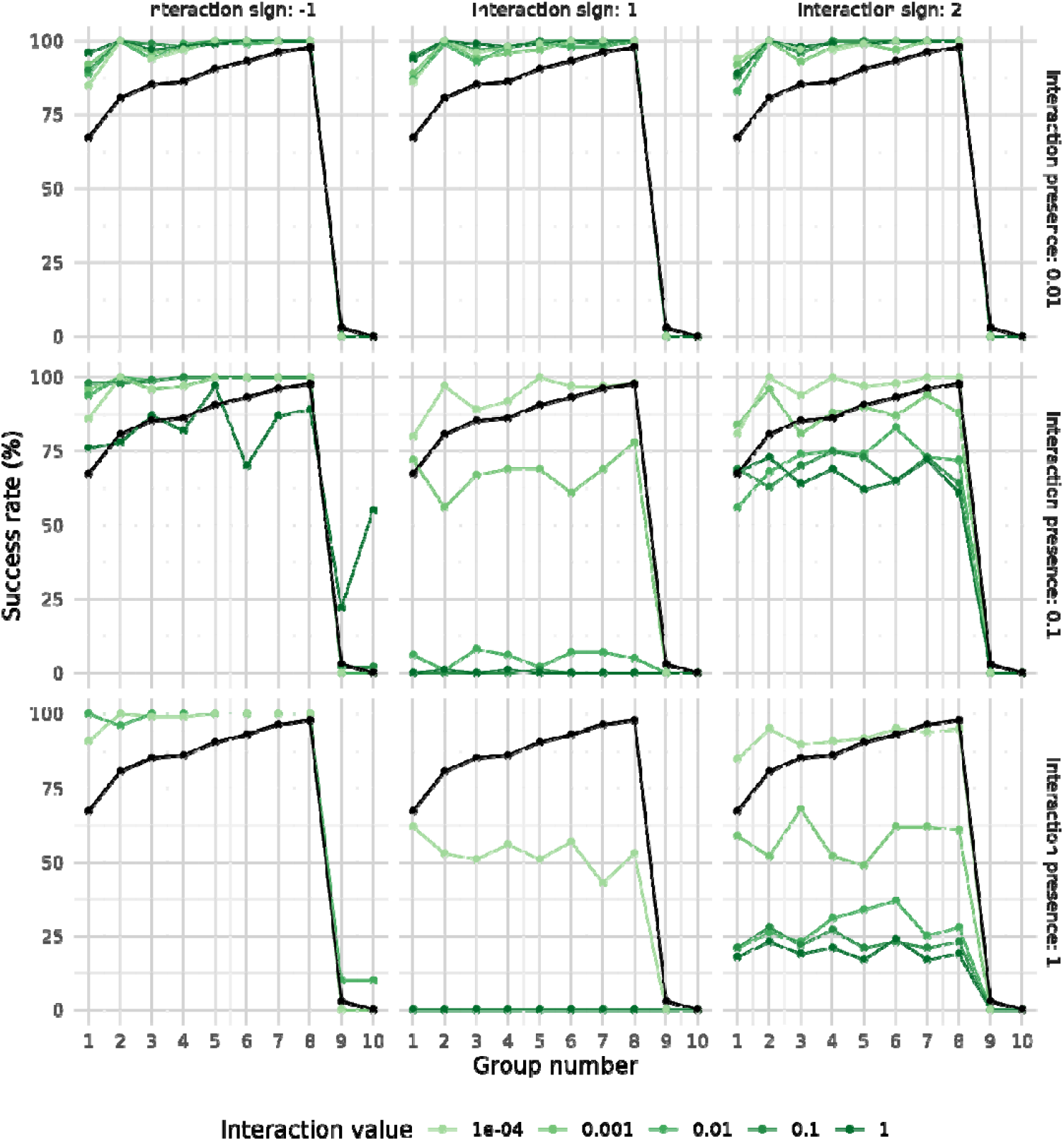
Success rate by functional group number in communities with 10 groups (heterogeneous group sizes, richness 100) after simulations under different interaction scenarios. Results without interactions are shown in black, while results under varying interaction strengths are shown in different colors, indicating both the presence and the sign of the interaction. A dilution factor of 0.1 was used in all simulations.

Additionally, while there was a clear positive correlation between group size and fixation in the simulations without interactions (with the exception of the two smallest groups, 9 and 10), the inclusion of interactions tended to reduce this correlation by leveling fixation rates across groups. However, the two smallest groups, being more vulnerable to extinction, generally did not benefit from negative interactions. Slight improvements in their fixation rates were observed only in two cases: weak negative interactions with 100% frequency (stronger interactions at this frequency were excluded due to growth suppression), and the strongest negative interactions with 10% frequency.

## DISCUSSION

Our simulations indicate that community size, dilution factor, and, to a lesser extent, initial community diversity, will significantly affect the success of the proposed strategy. There will be a delicate balance between the chosen dilution factor and the fixation or extinction of functional groups within the community, with marked differences between groups depending on their niche size. Furthermore, inter-group interactions will likely enhance the success of the approach when they are infrequent or weak (particularly if they are positive).

Based on our results, we conclude that while the successful implementation of the proposed approach for generating Minimal Microbiomes is feasible, it is constrained by several factors and will largely depend on the interplay between community size and dilution factor. However, even with an optimal balance between these parameters, functional groups with smaller niche sizes may still go extinct. Notably, the existence of inter-group interactions will likely increase the success of the approach when such interactions are either positive and infrequent, or negative but limited in frequency or intensity

Therefore, in the practical applications of this process to generate Minimal Microbiomes, perfect outcomes should not be expected; some functional loss must be assumed as part of the process. Alternatively, one could apply less stringent dilution factors and accept limited fixation among the more abundant functional groups. Nonetheless, in the former case, the missing minority functional groups could potentially be recovered through a second iteration of the process. This would involve reapplying the approach using the previously obtained minimal microbiome co-inoculated with a small proportion of the original community. In doing so, larger niches would already be occupied by the minimal microbiome, and the parameters of the second experiment could be tuned to favor recovery of smaller functional groups. Although this solution would effectively double the experimental workload, it may be the only viable strategy for obtaining a truly complete Minimal Microbiome. In any case, the proposed approach offers a powerful tool for simplifying experimental microbiomes, significantly enhancing our ability to isolate key populations, thus facilitating the construction of low-diversity consortia that remain functionally comprehensive and ecologically cohesive.

## Acknowledgements

This work was funded by the Spanish Ministry of Science and Innovation grant TED2021-130616B-I00 awarded to DAC.

## REFERENCES

1. Nemergut DR, Schmidt SK, Fukami T, O’Neill SP, Bilinski TM, Stanish LF, et al. Patterns and Processes of Microbial Community Assembly. Microbiology and Molecular Biology Reviews. 2013;77(3):342–56.

2. Chang C-Y, Vila JCC, Bender M, Li R, Mankowski MC, Bassette M, et al. Engineering complex communities by directed evolution. Nature Ecology & Evolution. 2021;5(7):1011–23.

3. Mueller UG, Sachs JL. Engineering Microbiomes to Improve Plant and Animal Health. Trends Microbiol. 2015;23(10):606–17.

4. Piccardi P, Vessman B, Mitri S. Toxicity drives facilitation between 4 bacterial species. Proc Natl Acad Sci U S A. 2019;116(32):15979–84.

5. Herrera Paredes S, Gao T, Law TF, Finkel OM, Mucyn T, Teixeira PJPL, et al. Design of synthetic bacterial communities for predictable plant phenotypes. PLoS biology. 2018;16(2):e2003962–e.

6. Jiang Y, Dong W, Xin F, Jiang M. Designing Synthetic Microbial Consortia for Biofuel Production. Trends Biotechnol. 2020;38(8):828–31.

7. Raaijmakers JM. The Minimal Rhizosphere Microbiome. In: Lugtenberg B, editor. Principles of Plant-Microbe Interactions: Microbes for Sustainable Agriculture. Cham: Springer International Publishing; 2015. p. 411–7.

8. Ridaura VK, Faith JJ, Rey FE, Cheng J, Duncan AE, Kau AL, et al. Gut microbiota from twins discordant for obesity modulate metabolism in mice. Science. 2013;341(6150):1241214.

9. Cordero OX, Datta MS. Microbial interactions and community assembly at microscales. Current Opinion in Microbiology. 2016;31:227–34.

10. Aguirre de Cárcer D. A conceptual framework for the phylogenetically constrained assembly of microbial communities. Microbiome. 2019;7(1):142.

11. Goldford JE, Lu N, Bajić D, Estrela S, Tikhonov M, Sanchez-Gorostiaga A, et al. Emergent simplicity in microbial community assembly. Science. 2018;361(6401):469–74.

12. Morella NM, Weng FC, Joubert PM, Metcalf CJE, Lindow S, Koskella B. Successive passaging of a plant-associated microbiome reveals robust habitat and host genotype-dependent selection. Proc Natl Acad Sci U S A. 2020;117(2):1148–59.

13. Sagova-Mareckova M, Omelka M, Kopecky J. The Golden Goal of Soil Management: Disease-Suppressive Soils. Phytopathology. 2023;113(4):741–52.

14. Wegmann U, Carvalho AL, Stocks M, Carding SR. Use of genetically modified bacteria for drug delivery in humans: Revisiting the safety aspect. Scientific Reports. 2017;7(1):2294.

15. Rocha I, Ma Y, Souza-Alonso P, Vosatka M, Freitas H, Oliveira RS. Seed Coating: A Tool for Delivering Beneficial Microbes to Agricultural Crops. Front Plant Sci. 2019;10(1357).

