## Supplementary for "Computational simulation of ecological drift for generating functional minimal microbiomes identifies key experimental and biotic factors influencing success"

^1^Microbial and Environmental Genomics Group, Departamento de Biología, Universidad Autónoma de Madrid, Madrid, Spain.

**Supplementary Figure** 1. Examples of abundances of four populations over time under three simulated scenarios: (A) without interactions, (B) with a negative interaction (P1 inhibits P3), and (C) with a positive interaction (P1 promotes the growth of P2). Solid lines represent the populations of Functional Group 1 (P1 and P2, carrying capacity 60%), while dashed lines represent the populations of Functional Group 2 (P3 and P4, carrying capacity 40%). Lines of different colors represent the abundance of each population over 10 successive dilution-growth cycles.

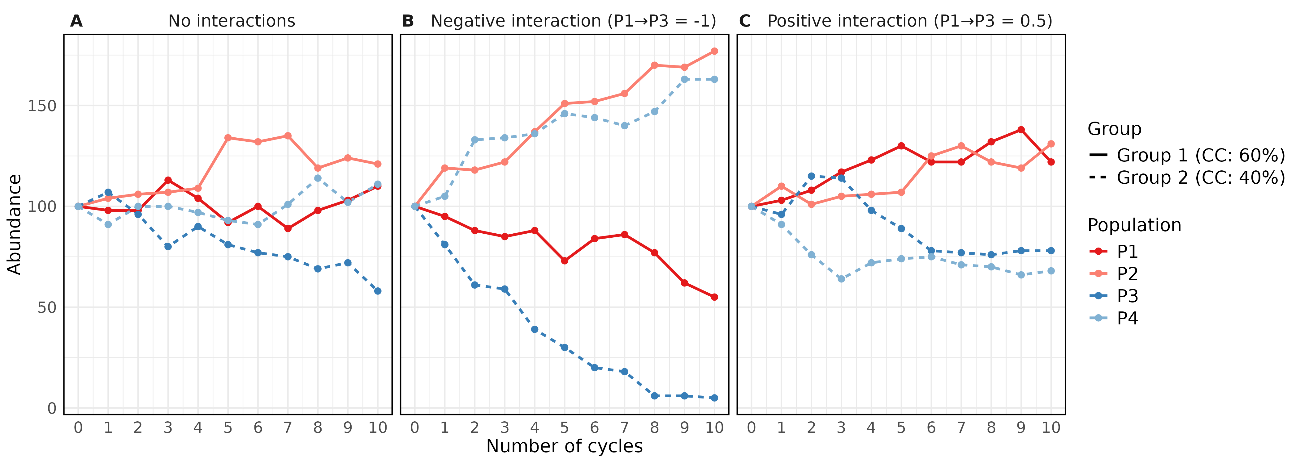

**Supplementary Figure 2**. Success patterns of dilution-growth processes according to the simulation parameters used. The Y axis indicates the dilution-growth cycle where success occurred, while the X axis orders the different simulated communities by dilution factor, distribution, community size, and richness. Each experiment is colored according to community type (same values of community size, abundance distribution, and richness); color intensity represents the dilution factor. Left; fixation threshold of 90%, Right; fixation threshold of 50%.

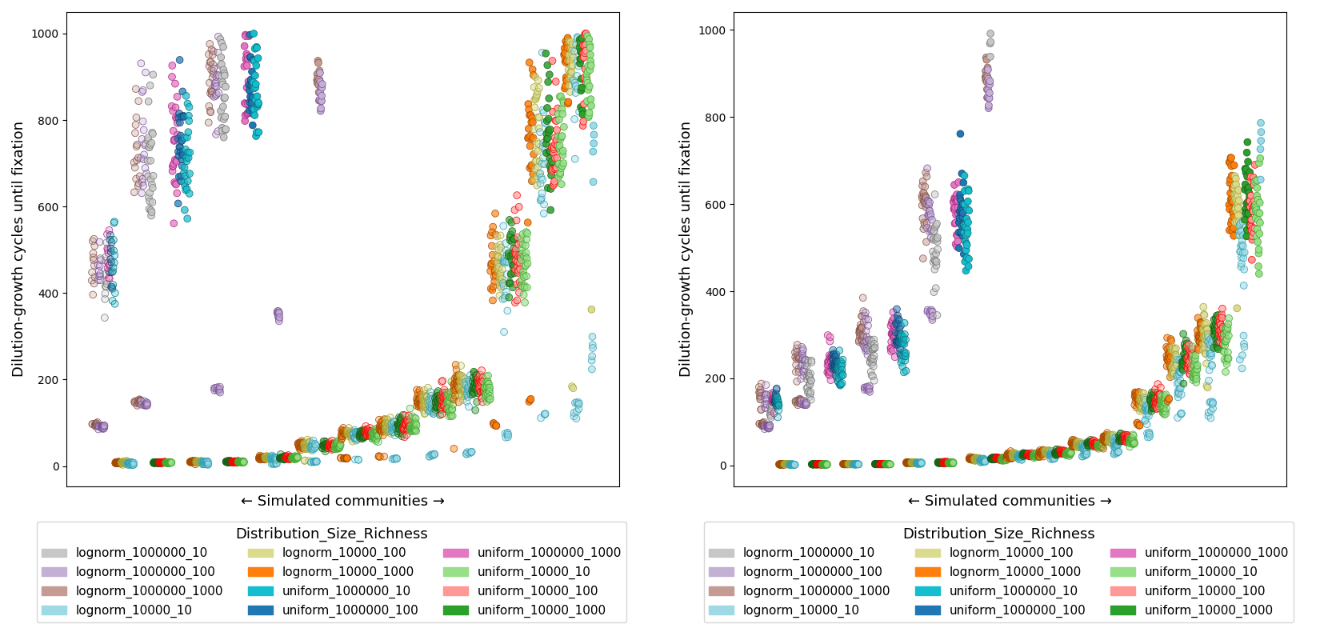

**Supplementary Figure 3**. Success rate for each of the 30 simulated communities with 10 functional groups, heterogeneous relative abundances, and a richness of 1000. Each plot corresponds to a different dilution factor. The Y axis indicates the proportion (from 0 to 1) of dilution-growth simulations in which total success occurs; that is, fixation in all functional groups. Each point represents one of the 30 communities. The missing point corresponds to a community that, after experiencing extinctions and a drop in total abundance, could not be sustained under the applied dilution factor.

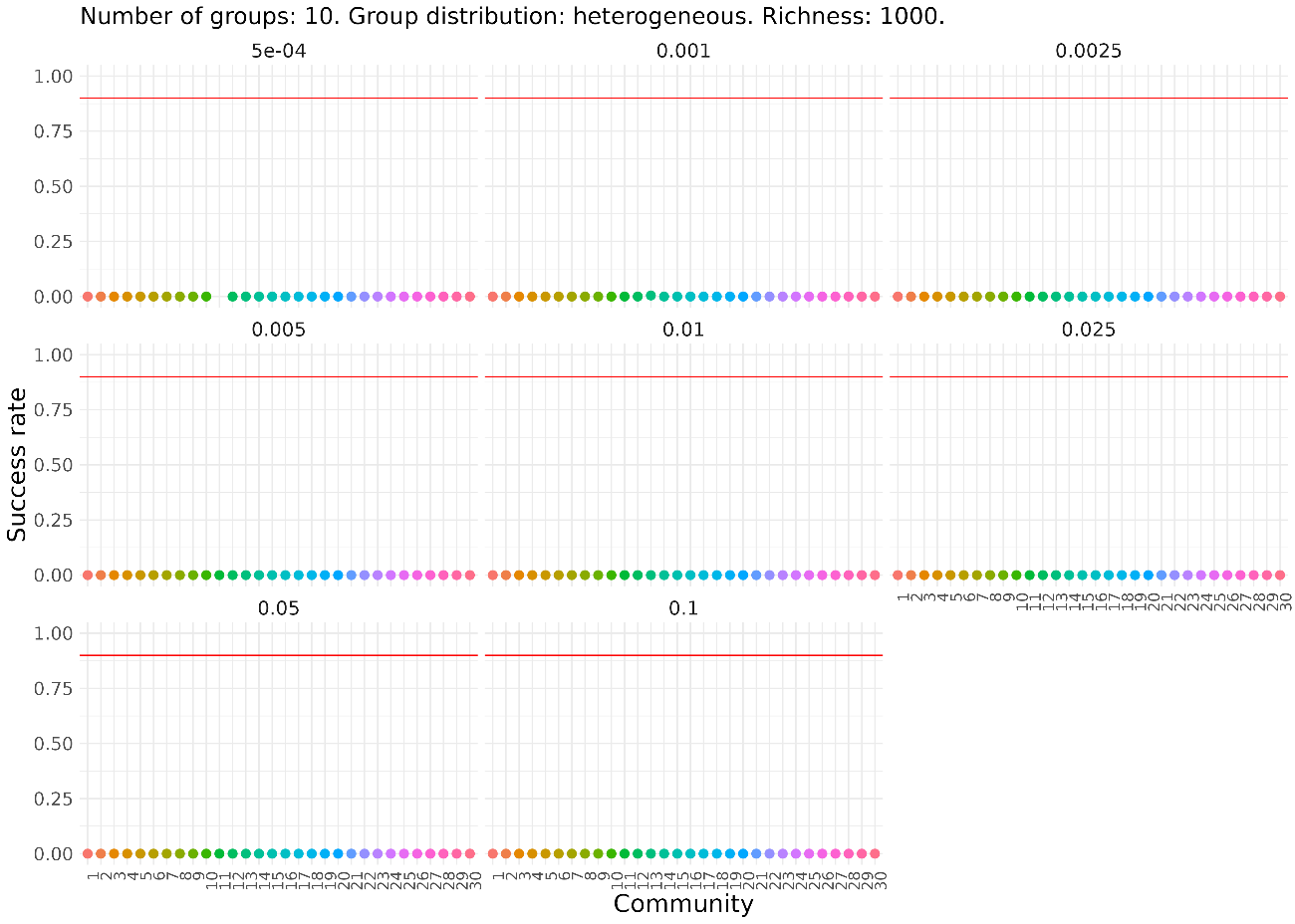

**Supplementary Figure 4**. Success rate for each of the 30 simulated communities with 10 functional groups, heterogeneous relative abundances, and a richness of 100. Each plot corresponds to a different dilution factor. The Y axis indicates the proportion (from 0 to 1) of dilution-growth simulations in which total success occurs; that is, fixation in all functional groups. Each point represents one of the 30 communities.

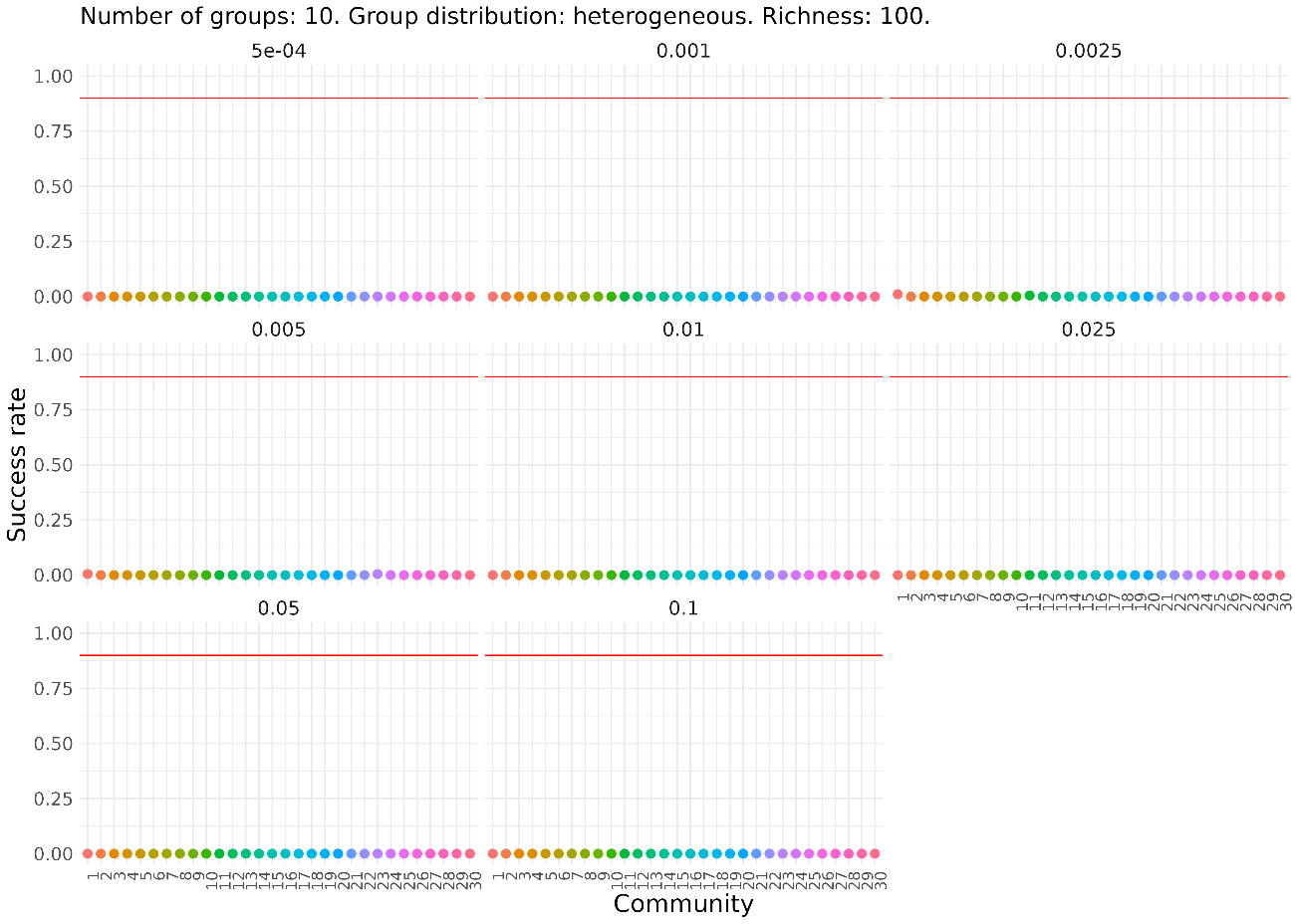

**Supplementary Figure 5**. Success rate for each of the 30 simulated communities with 3 functional groups, homogeneous relative abundances, and a richness of 1000. Each plot corresponds to a different dilution factor. The Y axis indicates the proportion (from 0 to 1) of dilution-growth simulations in which total success occurs; that is, fixation in all functional groups. Each point represents one of the 30 communities.

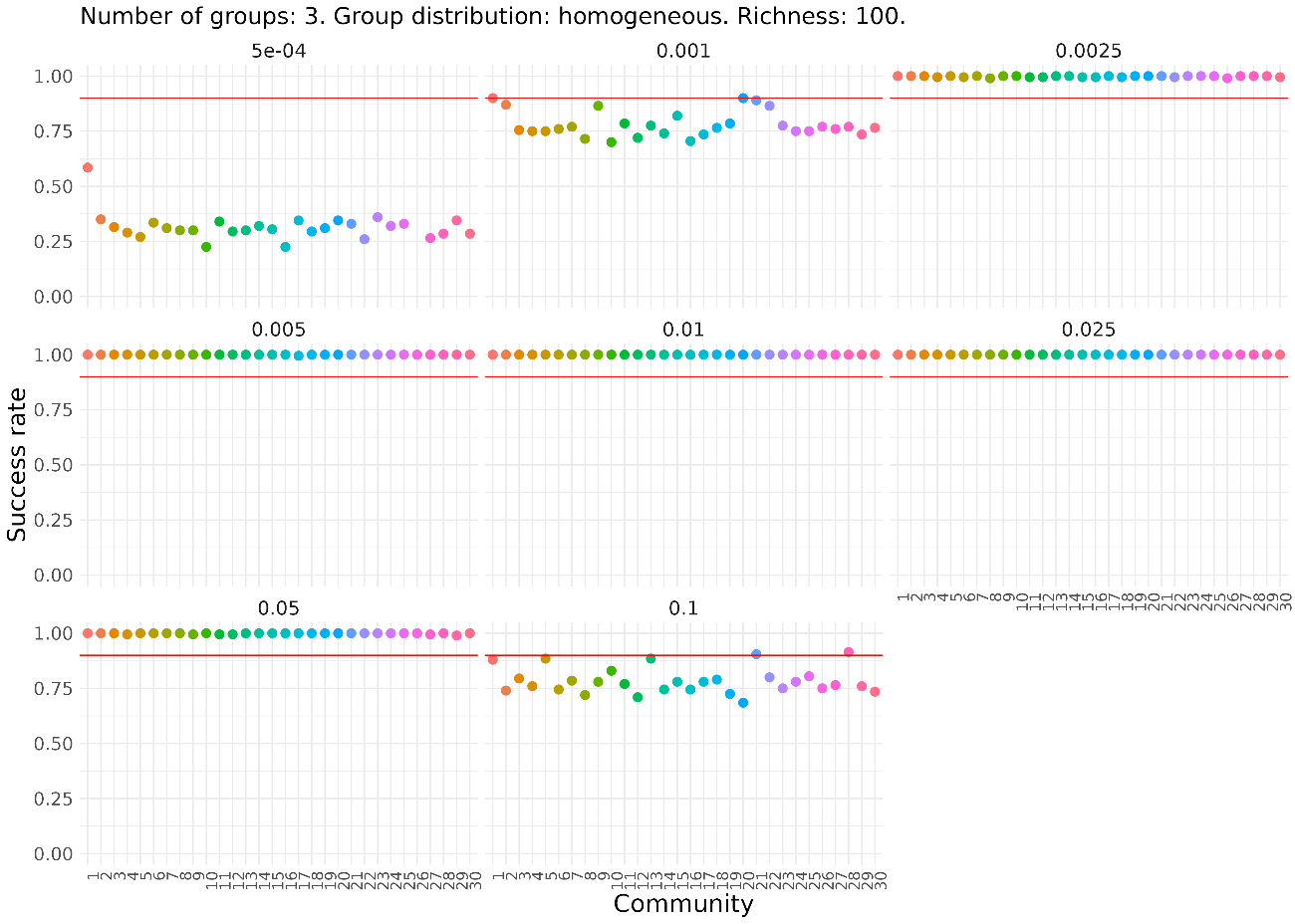

**Supplementary Figure 6**. Success rate for each of the 30 simulated communities with 3 functional groups, homogeneous relative abundances, and a richness of 100. Each plot corresponds to a different dilution factor. The Y axis indicates the proportion (from 0 to 1) of dilution-growth simulations in which total success occurs; that is, fixation in all functional groups. Each point represents one of the 30 communities.

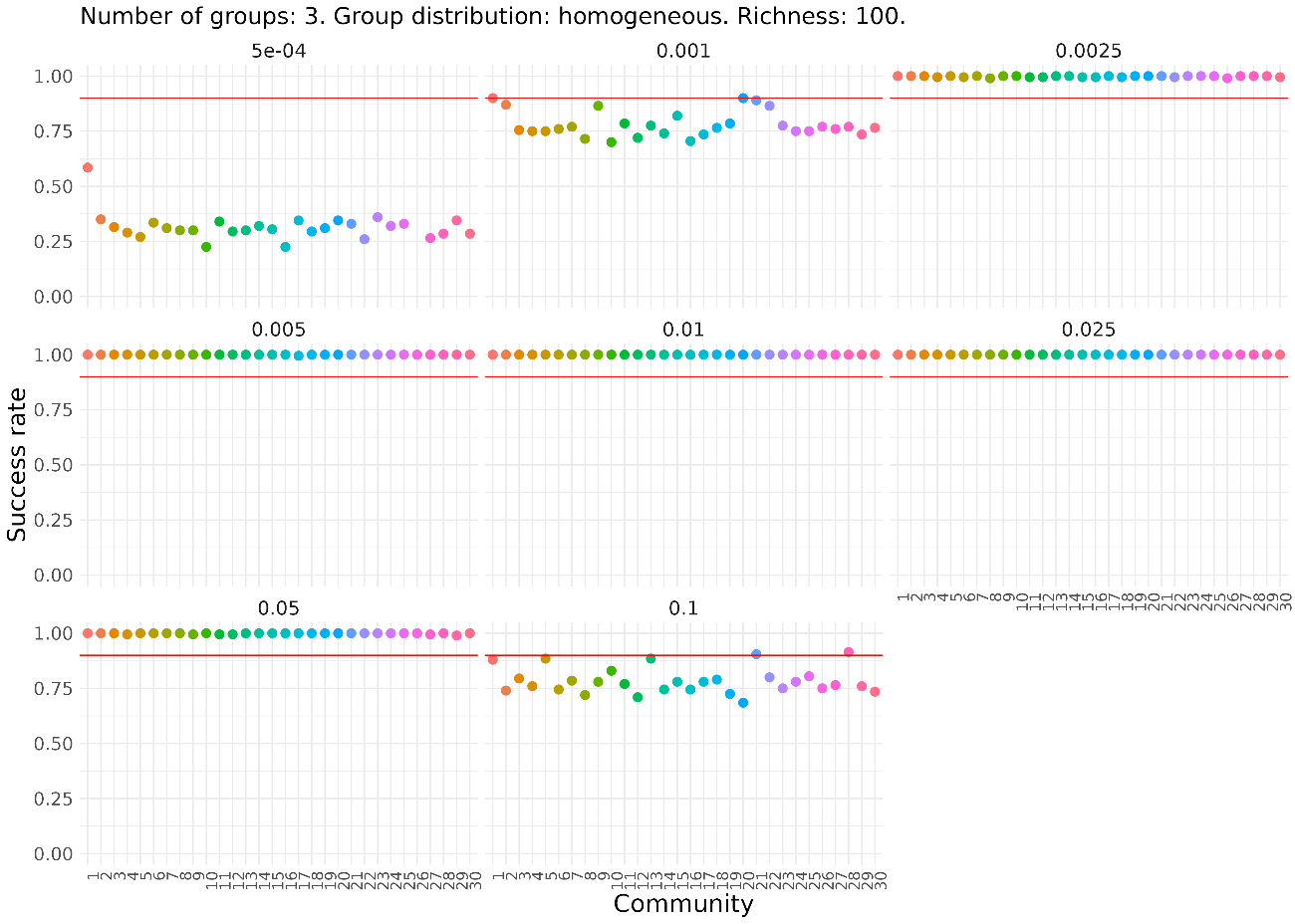

**Supplementary Figure 7**. Fixation and extinction rate by functional group in communities with homogeneous niche abundances. Results are shown for communities with a richness of 1000 and 3 or 10 groups (fixation in red and green, respectively, extinction in dark grey). Fixation for groups that exceed the 95% success threshold is highlighted in a darker shade. Bars represent the niche size associated with each functional group. Fixation is measured as the percentage of simulations in which at least one population from the group reaches fixation. Extinction is measured as the percentage of simulations in which no population from the group reaches fixation.

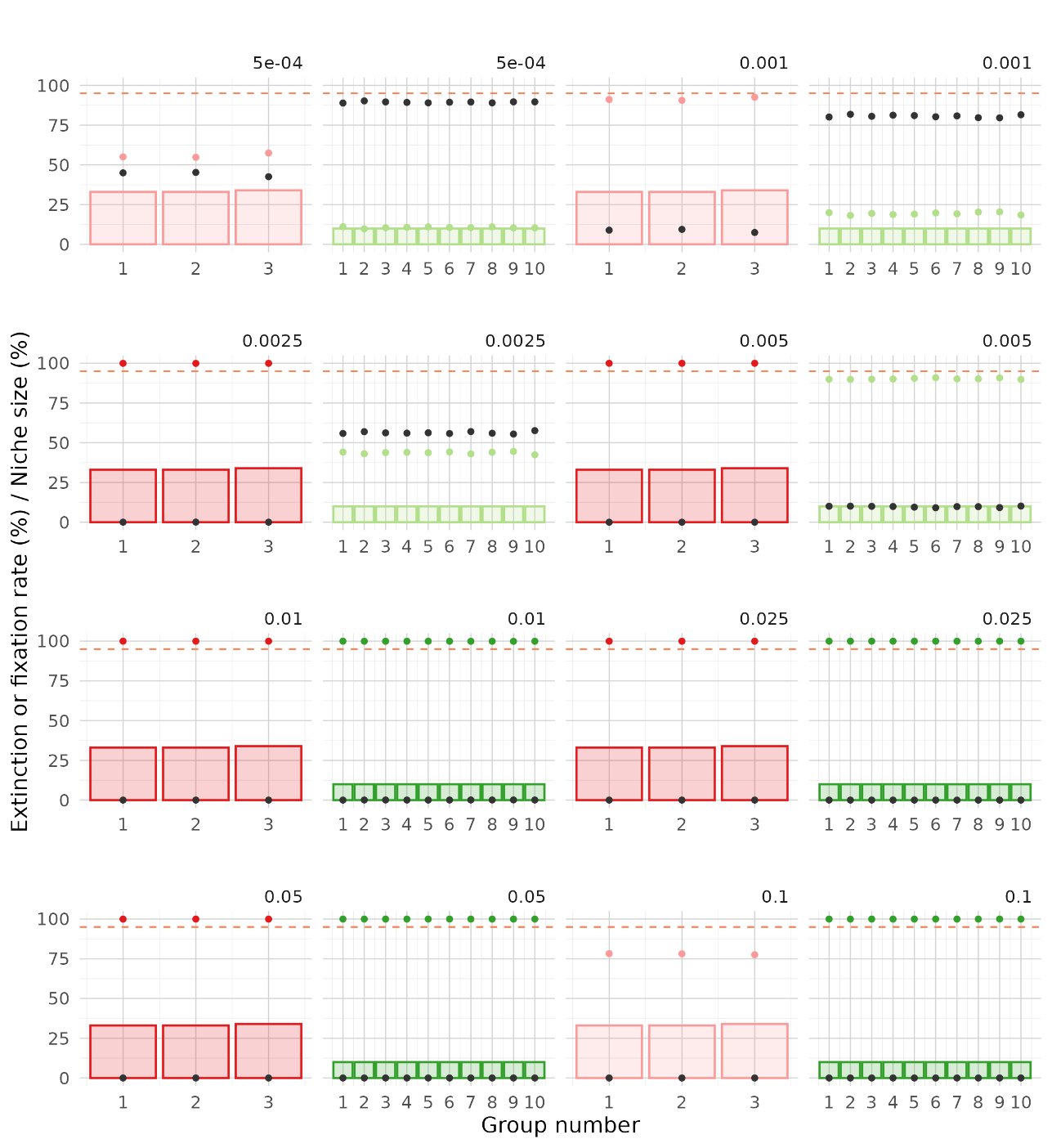

**Supplementary Figure 8**. Fixation and extinction rate by functional group in communities with heterogeneous niche abundances. Results are shown for communities with a richness of 1000 and 3 or 10 groups (fixation in red and green, respectively, extinction in dark grey). Fixation for groups that exceed the 95% success threshold is highlighted in a darker shade. Bars represent the niche size associated with each functional group. Fixation is measured as the percentage of simulations in which at least one population from the group reaches fixation. Extinction is measured as the percentage of simulations in which no population from the group reaches fixation.

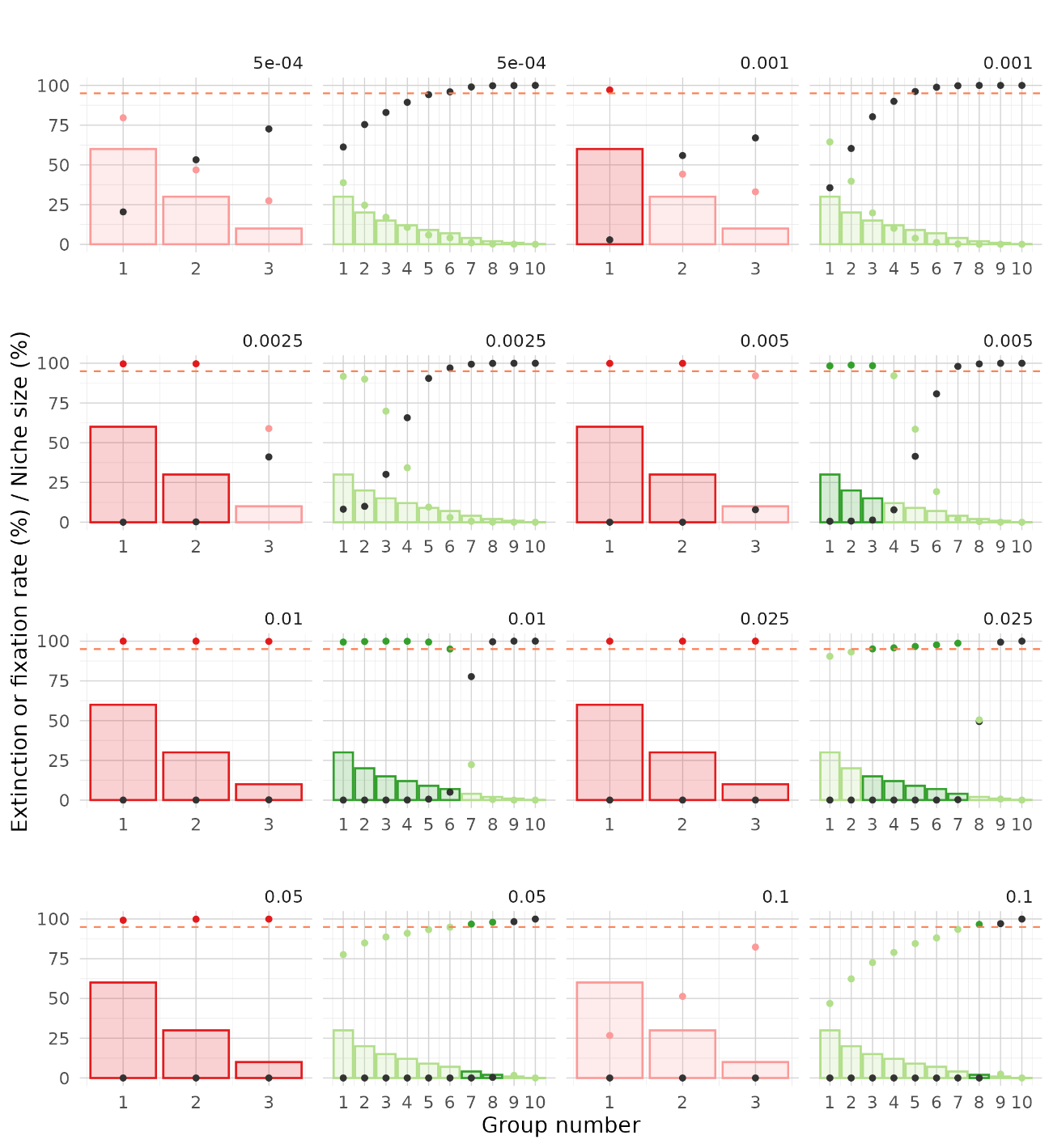

**Supplementary Table 1**. Mean decrease in accuracy for each variable used in the following models: (1) Pielou’s evenness only, (2) Shannon diversity only, (3) Gini index only, (4) Evenness and richness, (5) Shannon diversity and richness, (6) Gini index and richness. Success is defined using a fixation threshold of 50%.

| Model | Community size | Dilution factor | Pielou’s evenness | Shannon diversity | Gini index | Richness |
| --- | --- | --- | --- | --- | --- | --- |
| 1 | 0.576 | 0.509 | 0.033 | NA | NA | NA |
| 2 | 0.573 | 0.509 | NA | 0.038 | NA | NA |
| 3 | 0.575 | 0.509 | NA | NA | 0.032 | NA |
| 4 | 0.574 | 0.509 | 0.029 | NA | NA | 0.024 |
| 5 | 0.573 | 0.509 | NA | 0.039 | NA | 0.014 |
| 6 | 0.574 | 0.509 | NA | NA | 0.029 | 0.035 |

**Supplementary Table 2**. Mean decrease in accuracy for each variable used in the following models: (1) Pielou’s evenness only, (2) Shannon diversity only, (3) Gini index only, (4) Evenness and richness, (5) Shannon diversity and richness, (6) Gini index and richness. Success is defined using a fixation threshold of 90%.

| Model | Community size | Dilution factor | Pielou’s evenness | Shannon diversity | Gini index | Richness |
| --- | --- | --- | --- | --- | --- | --- |
| 1 | 0.540 | 0.52 | 0.072 | NA | NA | NA |
| 2 | 0.539 | 0.52 | N | 0.091 | NA | NA |
| 3 | 0.537 | 0.52 | NA | NA | 0.083 | NA |
| 4 | 0.539 | 0.52 | 0.073 | NA | NA | 0.056 |
| 5 | 0.539 | 0.52 | NA | 0.083 | NA | 0.064 |
| 6 | 0.539 | 0.52 | NA | NA | 0.081 | 0.052 |

**Supplementary Table 3**. R² values for each of the following models, defined according to the parameters they include: (1) Pielou’s evenness only, (2) Shannon diversity only, (3) Gini index only, (4) Evenness and richness, (5) Shannon diversity and richness, (6) Gini index and richness.

| Fixation threshold | 1 | 2 | 3 | 4 | 5 | 6 |
| --- | --- | --- | --- | --- | --- | --- |
| 50% | 0.990 | 0.991 | 0.990 | 0.991 | 0.991 | 0.991 |
| 90% | 0.929 | 0.936 | 0.934 | 0.937 | 0.937 | 0.93 |
